# Developing human fetal skin demonstrates a unique lymphocyte signature

**DOI:** 10.1101/2020.06.16.153593

**Authors:** Miqdad O. Dhariwala, Dhuvarakesh Karthikeyan, Kimberly S. Vasquez, Sepideh Farhat, Keyon Taravati, Elizabeth G. Leitner, Mariela Pauli, Margaret M. Lowe, Michael D. Rosenblum, Tiffany C. Scharschmidt

**Author notes:** Corresponding author Address: 1701 Divisadero Street, 3^rd^ Floor, San Francisco, CA 94115.

## Abstract

Lymphocytes in barrier tissues play critical roles in host defense and homeostasis. These cells take up residence in tissues during defined developmental windows, when they may demonstrate distinct phenotypes and functions. Here, we utilized mass and flow cytometry to elucidate early features of human skin immunity, demonstrating a unique fetal skin lymphocyte signature. While most conventional αβ T (Tconv) cells in fetal skin have a naïve, proliferative phenotype, a subset of CD4^+^ Tconv and CD8^+^ cells demonstrate memory-like features and a propensity for IFNγ production. Skin regulatory T cells dynamically accumulate over the second trimester in temporal and regional association with hair follicle development. These fetal skin Tregs demonstrate an effector memory phenotype while differing from their adult counterparts in expression of key effector molecules. Thus, we identify features of prenatal skin lymphocytes that may have key implications for understanding antigen and allergen encounters *in utero* and in infancy.

## Introduction

Each square centimeter of adult human skin contains approximately one million lymphocytes, comprised predominantly of CD4^+^ and CD8^+^ alpha beta (αβ) T cells^1^. These cells help defend us against cutaneous pathogens and malignancy and also facilitate key homeostatic tissue functions, such as wound healing and hair follicle cycling^2,3^. Conversely, they can play a central pathogenic role in common inflammatory and allergic skin diseases. As compared to our relatively nuanced understanding of lymphocytes in adult human skin, little is known about the phenotype and functional capacity of these cells during early life. Deciphering this initial immune landscape has the potential to critically inform our understanding of the role key lymphocyte populations play in normal human skin development as well as in immune-mediated skin diseases that begin early in life.

Human skin begins as a single-cell epithelium during embryogenesis and evolves over the first trimester of fetal life into a stratified epidermis with an overlying periderm. Epidermal differentiation and development of skin appendages occur from 15 to 20 weeks gestation, followed by acquisition of a functional stratum corneum between 20-24 weeks^4^. *In utero* maturation of skin architecture is accompanied by parallel seeding of the tissue by immune cells. Antigen presenting cells, found in the skin of 9-week-old embryos, are perhaps the first skin-resident population^5^. These are followed by T cells, which exit the fetal thymus around 11-14 weeks^6,7^, and are detectable in skin by the second trimester; *i.e.*, around 17-18 weeks^8,9^. To date, studies of human fetal tissues have largely employed immuno-histochemistry or flow cytometry to examine particular immune cell types of interest^8,9^. They have yet to provide a holistic understanding of the immune landscape in developing fetal skin or a deep investigation into the specific identities and phenotypes of fetal skin lymphocytes.

The fetal immune system is not merely a miniature version of that in adults. Rather, it exhibits cellular phenotypes and functions specifically adapted to the needs of the developing fetus and future neonate^10^. Perhaps most striking is a propensity towards adaptive immune tolerance, attributable to both lymphocyte-intrinsic and myeloid cell-dependent features^11-13^, which help limit excess inflammation to maternal, self or other *in utero* antigens. Recent studies of the fetal intestine have identified unique populations of cytokine-producing classical and innate-like effector T cells. These are thought to promote healthy gut development *in utero*, but may also modulate risk of inflammatory diseases in preterm neonates^11,12^.

Here we use a combination of flow and mass cytometry to elucidate the composition and phenotype of lymphocytes in human fetal skin with a focus on the second trimester as a particularly dynamic period. Our findings offer insight into unique features of fetal skin immunity, including the *in utero* presence of memory-like conventional T cells and an intimate relationship between accumulation of Tregs and skin morphogenesis. These findings may have important implications for cutaneous immune responses to self and foreign antigens *in utero* as well as in human infancy.

## Results

### Mass cytometry elucidates major immune cell subsets in human fetal skin

To obtain a broad understanding of the immune cells residing in human fetal skin, we performed mass cytometry (CyTOF) using a 22-antibody panel designed to include major lymphocyte and myeloid markers. 23 weeks gestation was chosen as a mature late second trimester fetal timepoint, and 5 torso skin samples were processed for CyTOF alongside 5 site-matched adult skin controls. Unbiased clustering of live-CD45^+^ cells was performed based on relative expression of the 22 markers and iterative analyses demonstrated that binning into 15 clusters captured the major phenotypic differences present (Fig 1a). Further analyses of the Uniform Manifold Approximation and Projection (UMAP) plots revealed clusters of immune cell populations, identities of which were assigned based on relative expression of key markers (Fig 1b). Most clusters were present in fetal and adult skin, however, a few were unique to fetal tissue.

**Figure 1:**
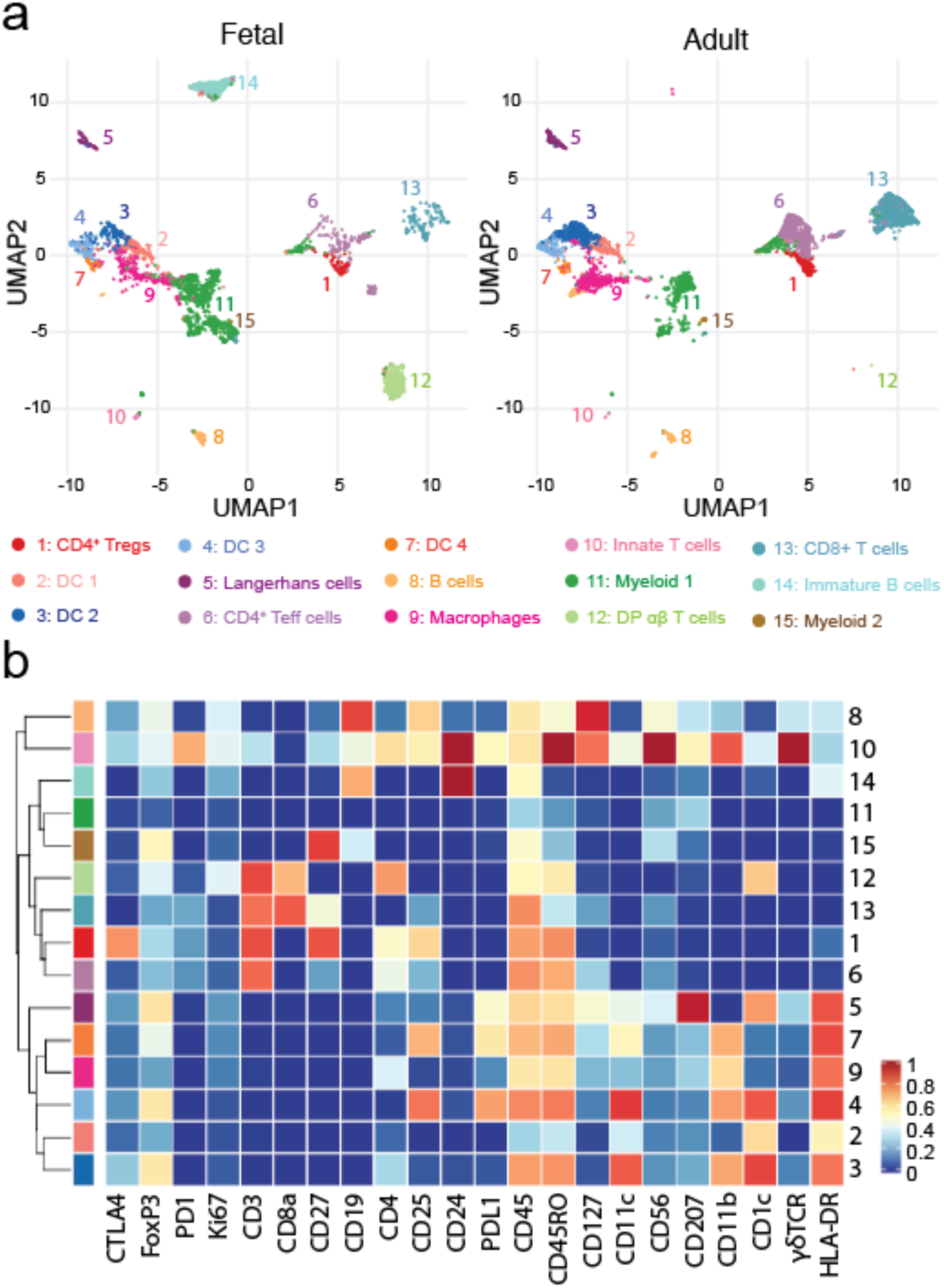
Mass cytometry reveals immune cell heterogeneity between fetal and adult skin. 23 week g.a. fetal torso skin along with healthy adult skin torso samples were analyzed in parallel for 22 markers using mass cytometry. (a) Uniform Manifold Approximation and Projection (UMAP) plots of viable CD45^+^ cells in fetal vs. adult skin, cells colored by cluster identity and plots annotated with cluster numbers and assigned identities. (b) Heatmap demonstrating relative expression by cluster of 22 markers in panel.

Fetal skin contained all major myeloid subtypes, including Langerhans cells (cluster 5), conventional type 1 dendritic cells (cDC1) (cluster 7), cDC2 (clusters 3 & 4), monocyte-derived DCs (cluster 2) and macrophages (cluster 9), which co-localized with their adult counterparts (Fig 1a). A subset of CD45^Lo^ cells (cluster 11) clustered near myeloid cells but was largely negative for markers in our panel and could not be definitively classified. In adult and fetal skin, CD19^+^HLA-DR^Int^ B cells constituted a very small population (cluster 8). In fetal skin, a second CD19^+^ subset was detected that co-expressed CD24^+^ (cluster 14), a marker typically seen on immature B cells and their precursors^13^. CD4^+^ (clusters 1 & 6) and CD8^+^ (cluster 13) T cells represented the bulk of lymphocytes present in fetal skin and were further expanded at the adult timepoint. Intriguingly, two of the five fetal skin samples contained a population of CD3^+^ cells co-expressing both CD4^+^ and CD8^+^ (cluster 12). However, flow cytometry performed on an additional 10 fetal skin samples failed to corroborate this as a consistent fetal skin T cell population (Fig S1). Non αβ T cells, i.e. γδ and/or NK T cells (cluster 10), constituted a very small fraction of lymphocytes at either age. Thus, human fetal skin contains shared but also unique immune cell subsets as compared to adult tissue. Among lymphocytes, CD4^+^ and CD8^+^ αβ T cells represent the largest lymphocyte population *in utero*.

### CD4^+^ and CD8^+^ Tconv cells in fetal skin largely demonstrate a naïve, proliferative phenotype

We next sought a more in-depth understanding of the types αβ T cells present in fetal versus adult skin. Re-clustering of CD4 single-positive cells across all samples revealed three major sub-populations (Fig 2a). Cluster C expressed high levels of Foxp3, CD25 and CTLA4 consistent with a regulatory T cell (Treg) phenotype, whereas clusters A and B were identifiable as two subsets of conventional CD4^+^ T cells (Tconv) (Supp Fig S2a-c). Whereas all adult CD4^+^ Tconv fell into cluster A, fetal Tconv were distributed across both clusters A and B (Fig 2b-c). This was corroborated by PCA analysis of CD4^+^ Tconv demonstrating fetal vs. adult segregation (Fig 2d). Skin CD8^+^ T cells displayed similar age-dependent clustering (Fig 2e-f).

**Figure 2:**
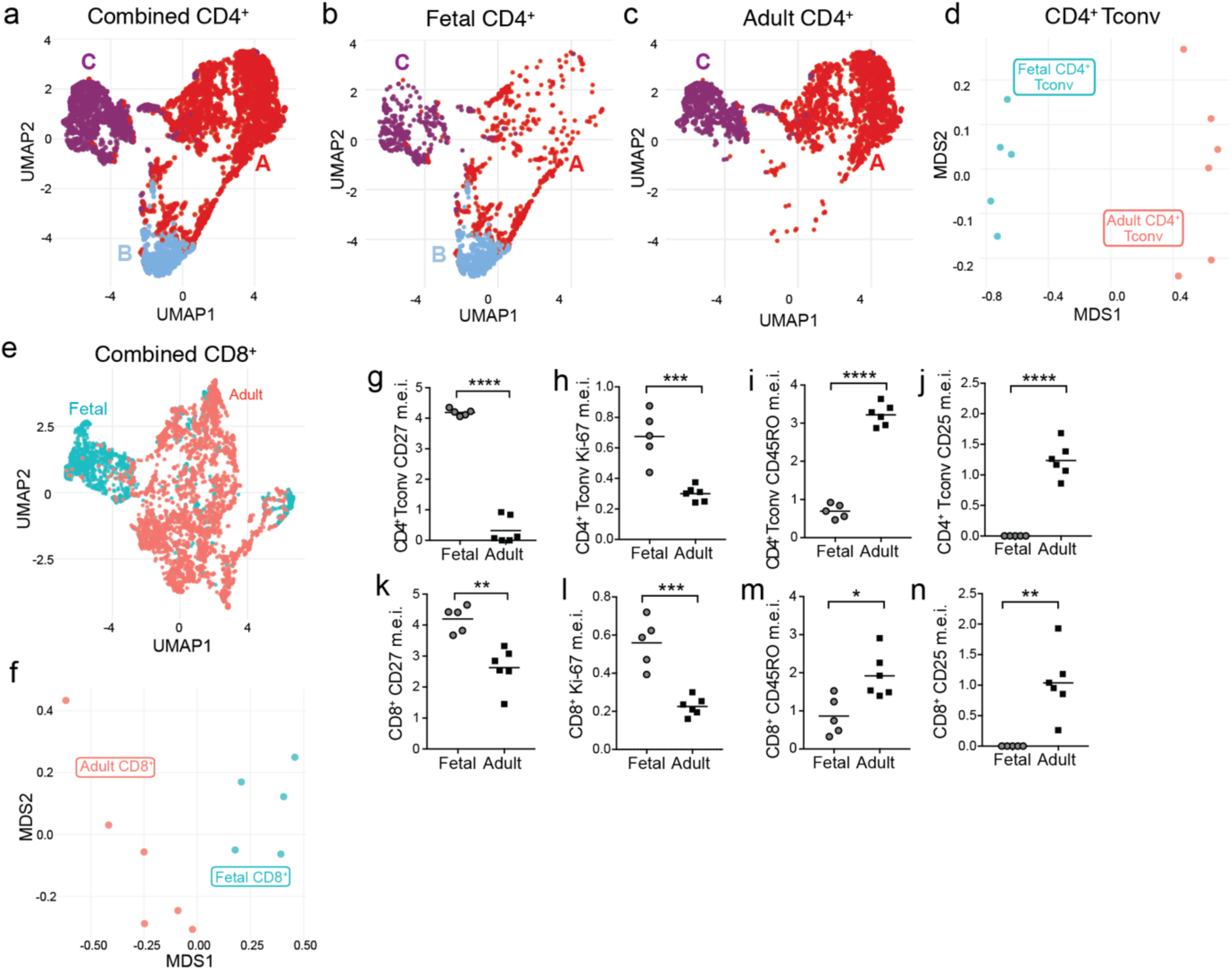
Conventional αβ T cells in fetal skin largely demonstrate a naïve, proliferative phenotype. (a) UMAP plots of CD4^+^ T cells from fetal and adult skin, combined. Cells are labelled and colored by cluster, with clusters A & B constituting conventional CD4^+^ T cells and cluster C representing Tregs. Analogous UMAP plots containing only (b) fetal or (c) adult CD4^+^ T cells. (d) Principal Component Analysis (PCA) plot demonstrating the distribution of conventional CD4^+^ T cells from each individual sample by age. (e) UMAP and (f) PCA plot demonstrating fetal and adult CD8^+^ T cell clustering by age. (g-n) Median expression intensity (m.e.i.) of (g, k) CD27, (h, l) Ki-67, (i, m) CD45RO and (j, n) CD25 for fetal vs adult CD4^+^ Tconv and CD8^+^ T cells as revealed by mass cytometric analyses. Each point in panels d, f, g-n represents data from an individual donor.

To understand the key drivers of this age-dependent clustering we calculated the mean expression intensity of markers in our panel on fetal versus adult CD4^+^ Tconv (i.e. those falling into clusters A & B above) and CD8^+^ cells. CD27, which is expressed often on naïve T cells^14^, and Ki-67, a marker of recent cell cycling, were elevated on fetal CD8^+^ and CD4^+^ Tconv (Fig 2g-h, k-l & S2d-e). CD3 and Foxp3 were likewise slightly enriched on fetal cells (Fig S2h-k). By comparison, CD45RO and CD25, markers of memory and activation, were enriched on adult skin CD4^+^ Tconv and CD8^+^ (Fig 2i-j, m-n & S2f-g). Thus, CD4^+^ Tconv and CD8^+^ T cells in fetal skin largely express markers indicative of a proliferative, naïve status.

### A subset of fetal skin CD4^+^ and CD8^+^ Tconv are CD45RO^+^ and demonstrate enriched capacity for IFNγ production

Notably, our CyTOF analysis revealed a subset of fetal CD4^+^ T cells that fell into cluster A alongside adult CD4^+^ Tconv (Fig 2b). These cells expressed relatively more CD45RO and CD25 and relatively less CD27 and Ki-67, suggestive of a memory phenotype (Fig S2d-g). To corroborate this observation and better quantify the proportion of fetal CD4^+^ Tconv and CD8^+^ cells with a naïve versus memory phenotype, we performed flow cytometry on a larger number of fetal skin samples including staining for CD45RA and CD45RO, markers respectively found on antigen-naïve and antigen-experienced lymphocytes. Strikingly, up to 30% of CD4^+^ Tconv and slightly fewer CD8^+^ T cells in fetal skin were CD45RO^+^ (Figs 3a-c).

**Figure 3:**
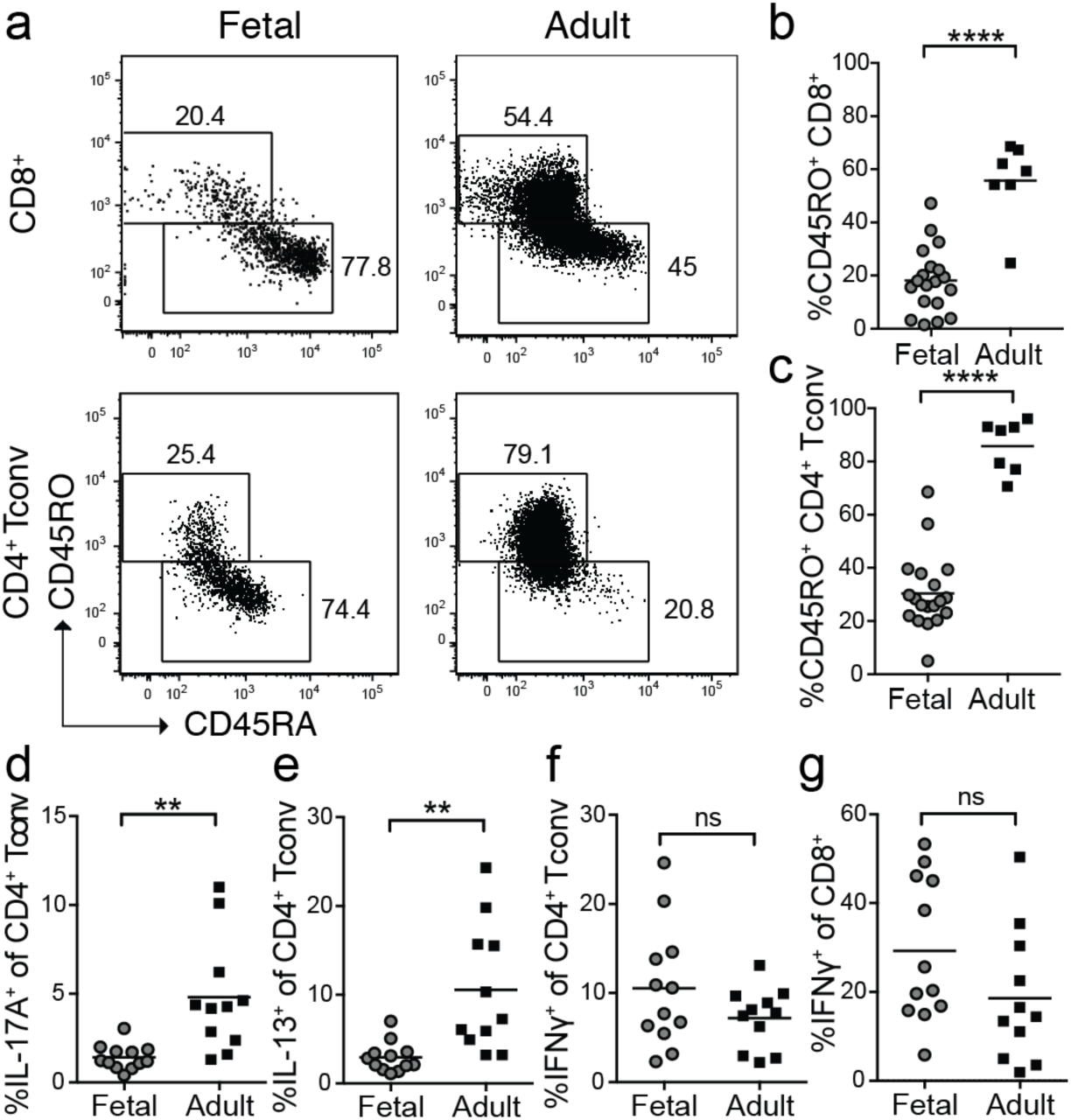
Subsets of fetal skin CD4^+^ Tconv and CD8^+^ T cells display a memory phenotype and demonstrate capacity for IFNγ production. Cells were isolated from second trimester fetal skin (scalp and/or torso) as well as adult (torso) skin and analyzed by flow cytometry. (a) Representative plots demonstrating CD45RO expression by fetal CD8^+^ T cells (gated on live CD3^+^CD8^+^CD4^neg^) and CD4^+^ Tconv (gated on live CD3^+^CD4^+^CD8^neg^Foxp3^neg^CD25^lo^). Percentage of CD45RO^+^ (b) CD8^+^ T cells and (c) CD4^+^ Tconv in fetal vs. adult skin. Percentage of CD4^+^ Tconv producing (d) IL-17A, (e) IL-13, and (f) IFNγ after PMA/ionomycin re-stimulation, and (g) percentage of IFNγ−producing CD8^+^ T cells. Each point in b-g represents data from an individual tissue sample; for some fetal samples data from scalp and torso skin from the same fetal donor are included as separate points.

To further characterize the functional potential of fetal skin T cells, we performed *ex vivo* re-stimulation with PMA and ionomycin followed by intracellular cytokine staining. TNFα is produced preferentially by memory as compared to naïve T cells^18,19^. Consistent with their largely naïve status, CD4^+^ Tconv and CD8^+^ T cells in fetal skin expressed substantially less TNFα than those in adult skin (Fig S3a & S3b). Production of IL-22, IL-17a and IL-13 were also reduced in fetal versus adult skin CD4^+^ Tconv (Figs 3d, 3e & S3c). In contrast, fetal lymphocyte production of IFNγ was equivalent to adult levels in CD4^+^ Tconv and trended higher in fetal CD8^+^ T cells (Fig 3f & 3g). To try to account and control for inherent differences in the type of Tconv cells found in fetal versus adult skin, *i.e.* frequency of CD45RO^+^ cells, we normalized production of each cytokine to that of TNFα on a per sample basis. This revealed enriched IFNγ production among fetal versus adult skin T cells (Figs S3d & S3e), whereas production of IL-13, IL-17 and IL-22 was equivalent or reduced (Figs S3f-h). Thus, human fetal skin contains a subset of conventional T cells that demonstrate a memory-like phenotype and propensity for IFNγ production.

### Fetal skin Tregs display properties of effector memory Tregs

The fetal immune system is uniquely poised for immune regulation^15^. We therefore, returned to our CyTOF analysis to carefully examine the phenotype of Tregs in fetal skin. As noted earlier, Tregs from both fetal and adult skin fell into CD4^+^ cluster C (Figure 2a-c; Fig S2a-c). Although skin Tregs also segregated by age upon PCA analysis, age-dependent differences were strikingly less than for CD4^+^ Tconv (Fig 4a). Dissecting this further, we found several markers distinguishing fetal versus adult skin Tregs. Akin to other cell types, Tregs in fetal skin expressed higher levels of Ki-67, CD27 and CD3 (Fig S4a-c). CD45RO expression was high among fetal Tregs versus CD4^+^ T conv cells but lagged slightly behind that of adult Tregs (Fig S4d). Notably, fetal skin Tregs demonstrated higher levels of the lineage-defining transcription factor, Foxp3 (Fig 4b) and equivalent amounts of the high affinity IL-2 receptor alpha chain, CD25 (Fig 4c) as compared to adult skin Tregs. With regard to other important Treg activation markers, fetal skin Tregs demonstrated a non-significant trend towards higher PD-1 levels (Fig 4d), but lower amounts of total CTLA4 (Fig 4e). Thus, while differing somewhat in their relative expression of certain markers, Tregs in fetal skin largely demonstrate an effector memory Treg phenotype, akin to Tregs in adult skin.

**Figure 4:**
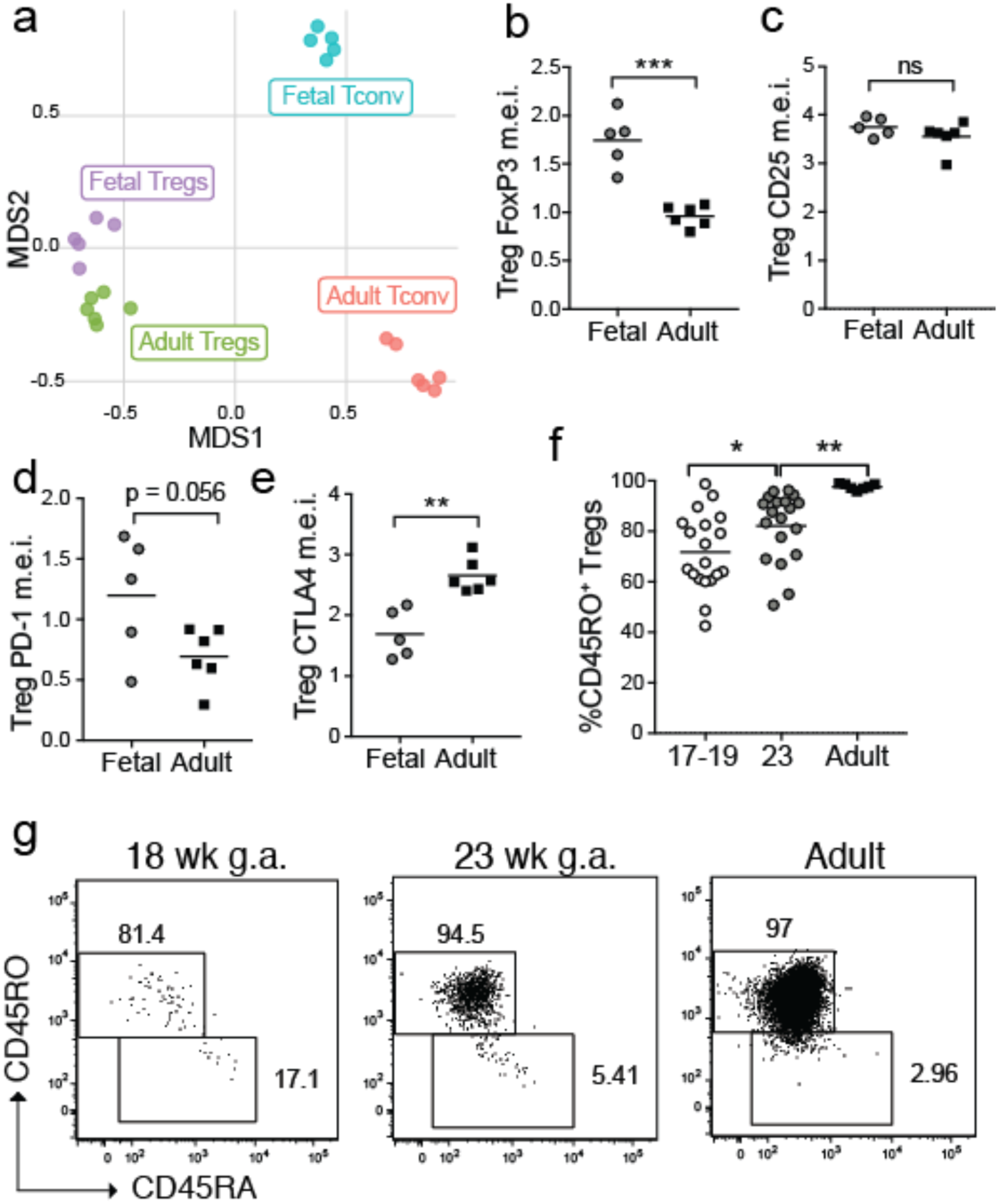
Tregs in human fetal skin demonstrate an effector memory phenotype. 23 week g.a. fetal torso skin and healthy adult torso skin samples were analyzed for 22 markers using mass cytometry. (a) PCA plot of Tregs (cluster C) and CD4^+^ Tconv (clusters A & B) from 23 week fetal skin vs. adult skin based on relative expression of key markers as assessed by mass cytometry. (b-e) Graphs showing median expression intensity (m.e.i.) of (b) Foxp3, (c) CD25, (d) PD-1, and (e) CTLA4 by mass cytometry on fetal vs. adult skin Tregs. (f-g) Cells were isolated from 17 to 23 week g.a. fetal skin (scalp and/or torso) as well as adult (torso) skin and analyzed by flow cytometry. (f) Percentage of CD45RO^+^ Tregs in skin by age and (g) accompanying representative flow plots. Each point in a-e represents data from an individual donor. Points in f point represent data from an individual tissue sample; for some fetal samples data from scalp and torso skin from the same fetal donor are included as separate points.

Tregs leaving the fetal thymus are CD45RO^neg16^. Thus, we hypothesized these cells were acquiring their memory phenotype in skin following exposure to self-antigen. To examine this possibility, we performed flow cytometry on additional samples, now including earlier second trimester fetal skin for comparison. In contrast to CD4^+^ Tconv and CD8^+^ cells which demonstrated a consistently low percentage of CD45RO^+^ cells independent of fetal age (Fig S4e-f), the proportion of CD45RO^+^ Tregs progressively increased during the second trimester (Fig 4f-g). Thus, the population of fetal skin Tregs evolves over the second trimester with progressive acquisition of a memory phenotype.

### T cells accumulate in fetal skin during the second trimester via both continued thymic egress and local proliferation

Having delineated features of late second trimester fetal skin lymphocytes, we next sought to better define dynamics of their tissue accumulation. Although CD3^+^ T cells represented a comparatively small percentage of total events in fetal versus adult skin, they were readily identifiable by flow cytometry in fetal samples as young as 17 weeks and substantially expanded by 23 weeks (Fig 5a and S5a). While the proportion of CD4^+^ T cells within the CD3 T cell compartment was stable across both fetal and adult time points (Fig 5b and S5b), the average percentage of Foxp3^+^ regulatory T cells (Tregs) among CD4^+^ cells doubled between 17 and 23 weeks gestation and was higher at this later fetal timepoint than in adult skin (Fig 5c and Fig S5c).

**Figure 5:**
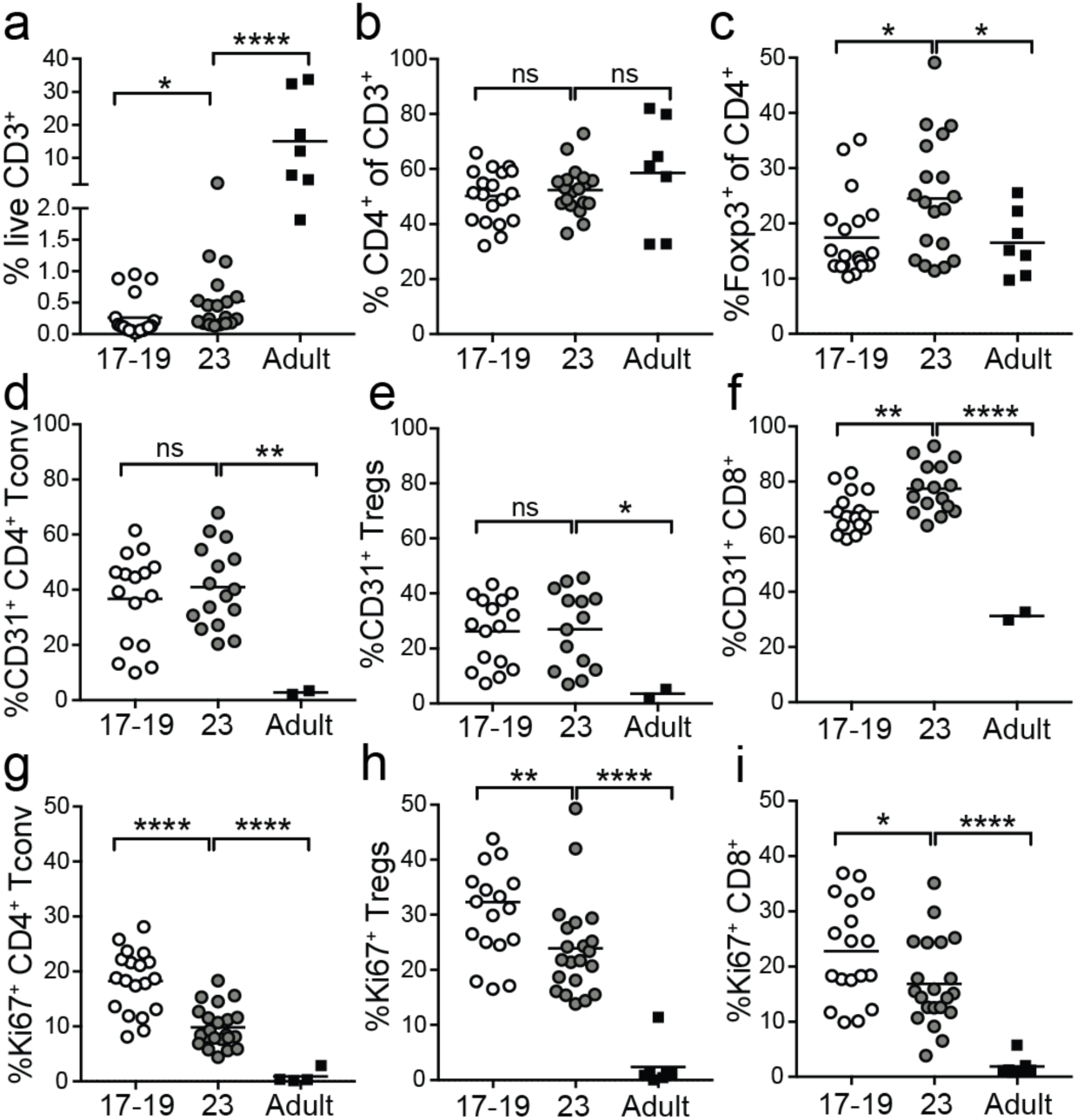
Lymphocytes progressively accumulate in fetal skin via a combination of thymic egress and local proliferation. Cells were isolated from 17 to 23 week g.a. fetal skin (scalp and/or torso) as well as adult (torso) skin and analyzed by flow cytometry. (a) Percentage of live CD3^+^ lymphocytes, (b) percentage of CD4^+^ T cells, and (c) percentage of Foxp3^+^ Tregs by age. (d) Percentage of CD31^+^ CD4^+^ Tconv cells, (e) Tregs, and (f) CD8^+^ T cells by age. (g) Percentage Ki-67^+^ CD4^+^ Tconv cells, (h) Tregs, and (i) CD8^+^ T cells by age. Points represent data from an individual tissue sample; for some fetal samples data from scalp and torso skin from the same fetal donor are included as separate points.

To elucidate the contribution of ongoing cellular influx on progressive T cell accumulation in fetal skin, we examined expression of CD31, a cell surface adhesion molecule found in high levels on recent thymic emigrants^17^. As compared to adult skin, CD31^+^ T cells were substantially enriched in fetal skin at both 17-18 and 23 weeks (Fig S5d). CD31 was expressed by 20 to 60 percent of fetal skin CD4^+^ cells, in comparison to less than 5% of adult CD4^+^ T cells (Fig 5d-e). As reported previously, CD8^+^ cells demonstrated comparative higher CD31 expression at all fetal ages relative to adult (Fig 5f)^18^. Of note, CD31 expression was significantly higher on CD45RO^neg^ versus CD45RO^+^ CD4^+^ Tconv and CD8^+^ cells in fetal skin, but CD45RO status did not correlate with CD31 expression among fetal skin Tregs (Fig S5f-h).

To determine the extent to which *in situ* proliferation also contributes to increasing numbers of T cells in fetal skin, we examined Ki-67 expression by flow cytometry (Fig S5e). Corroborating our CyTOF data, Ki-67 was expressed by less than 5 percent of adult skin T cells. In contrast, it was found on 15 to 40 percent of T cells in fetal skin at 17-18 weeks and slightly fewer at 23 weeks (Fig 5g-i). Whereas CD45RO expression was associated with only slightly higher Ki-67 levels among fetal skin CD4^+^ Tconv and CD8^+^ cells, CD45RO^+^ fetal skin Tregs expressed twice as much Ki-67 compared to their CD45RO^neg^ counterparts (Fig S5i-k). The proportion of Ki-67^+^ cells among skin Tregs was also higher at both fetal time points as compared to CD4^+^ Tconv or CD8^+^ T cells (Fig 5h). Thus, while recent thymic emigrants constitute a substantial fraction of T cells in second trimester fetal skin, cell proliferation also contributes to their progressive increase during this period, especially among CD45RO^+^ cells.

### Enrichment in fetal skin Tregs coincides with initial hair follicle morphogenesis

The observation that fetal skin Tregs increase in percentage during the second trimester (Fig 5c) and display unique dynamics of CD31 and Ki-67 expression (Fig 5d-i) prompted us to explore the role of tissue-intrinsic factors in their progressive accumulation. Because Tregs in adult human skin localize preferentially around hair follicles and are more abundant in hair-dense skin sites^17,18^, we investigated the relationship between the appearance of fetal Tregs and hair follicle development. We first examined histologic sections from our fetal skin torso samples and scored them by stage of hair follicle development^19^. Consistent with published reports^20^, we found that hair follicle development progressed significantly between 20 and 23 weeks gestation (Fig 6a & 6b), which coincided with the increased percentage of Tregs (Fig 6c). Next, we took advantage of the fact that fetal hair follicles mature in a cephalocaudal pattern^21^. Using paired skin samples from the scalp and torso of fetuses aged 20 weeks or less, when torso hair follicles are still relatively immature versus those on the scalp, we assessed the density of Tregs irrespective of fetal age. Indeed, for a given fetus, Treg percentages were higher in scalp as compared to torso skin (Fig 6d). These results demonstrate that skin Tregs increase contemporaneously with fetal hair follicle development *in utero*.

**Figure 6:**
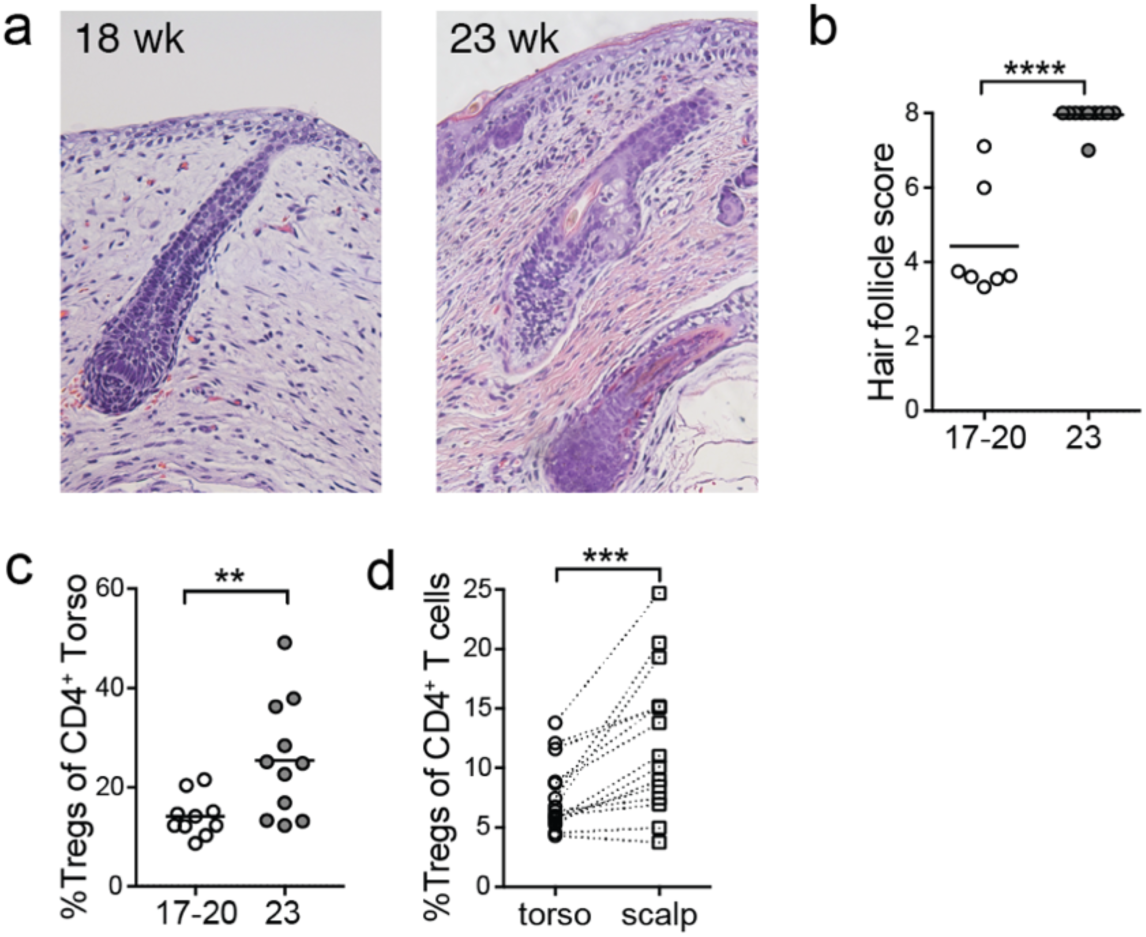
Treg accumulation in human fetal skin coincides spatiotemporally with second trimester hair follicle development. Fetal skin samples ranging from 17-23 weeks g.a. were processed for histology. (a) Representative images of tissue sections stained with hematoxylin and eosin from 18 and 23 weeks g.a. skin. (b) Hair follicles were scored and graphed with each data point representing cumulative scores from an individual donor. (c) Percentage Tregs in fetal skin at gestational age 17-20 versus 23 weeks as assessed by flow cytometry. (d) Percentage Tregs in paired torso versus scalp skin from fetal donors of less than 20 weeks g.a. Each matched set of data points originates from a different fetal skin donor.

## Discussion

Here we provide a holistic view of the cellular immune composition of human fetal skin, which already contains the vast majority of cell subtypes found in adult skin. Additionally, some unique, immature lymphocyte subsets, i.e. CD24^+^CD19^+^ B cells and CD4^+^CD8^+^CD3^+^ T cells, are sporadically detected in some but not all fetal samples. Conventional Foxp3^neg^CD4^+^ and CD8^+^ T cells in fetal skin are largely distinguishable from their adult counterparts by high expression of Ki-67 and CD27 and low expression of CD25 and CD45RO, indicative of a naïve phenotype. Some fetal Tconv cells, however, are positive for CD45RO and display preferential capacity for IFNγ production, suggesting the presence of a memory Tconv cell subset in fetal skin. Although differing in expression of certain effector molecules, fetal skin Tregs are quite similar to their adult counterparts and express markers identifying them as effector memory Tregs. All fetal skin lymphocytes, but especially CD45RO^neg^ Tconv are marked by elevated CD31 expression suggesting enrichment for recent thymic emigrants. Local proliferation suggested by Ki-67 is preferentially seen among CD45RO^+^ cells, in particular Tregs. Progressive accumulation of Tregs in fetal skin occurs contemporaneously with hair follicle development, and preferential accumulation occurs at sites of hair follicle density within a given fetus, suggesting an early role for hair follicle biology in skin Treg residency. Together these results illuminate key aspects of the immune landscape in human fetal skin, which have the potential to significantly inform our understanding of late prenatal and early postnatal immune responses as related to both health and disease.

In comparison to prior studies, we were able to leverage the capacity of CyTOF to broadly phenotype and simultaneously capture the relative abundance of both lymphoid and non-lymphoid immune cells within a tissue. Our 22-marker panel allowed us to identify most major immune cell types in adult skin and determine that these are also present during fetal development. We were intrigued to note that certain fetal samples also contained cell populations unique to this time period. A population of CD24^+^CD19^+^ cells likely represent B cell precursors, as described in human bone marrow^13^. We did not assess CD38 in our panel, but expression of this glycoprotein by CD24^+^CD19^+^ cells can indicate a transitional state with regulatory potential^22^. CD4^+^CD8^+^ T cells were not present in the majority of fetal samples we examined, but constituted a substantial lymphocyte subset in some. Double positive T cells represent an immature precursor state during lymphocyte development in the thymus. They have also been seen peripherally in certain immunologic diseases, including the skin of patients with atopic dermatitis, systemic sclerosis and graft versus host disease, where they are thought to potentially have a regulatory role via production of IL-10 or TGF-β^23^. In the context of fetal development, we speculate that these may represent immature cells that transiently or sporadically escape to the periphery where they either die or further differentiate into the subsets we see in mature skin. Further work is needed to explore whether these cells are present in other non-lymphoid fetal tissues and at what point during development they “disappear”. Prior studies have illuminated functional differences in fetal versus adult skin dendritic cells^24^. While not the focus of this work, a more comprehensive examination of different skin myeloid cell subsets by age is clearly warranted.

In these studies, we chose to more carefully elucidate the phenotype of lymphocytes, specifically T cells, residing in human fetal skin. Among CD4^+^ Tconv and CD8^+^ T cells, wide segregation by age was primarily driven by differential expression of Ki-67, CD45RO, CD25, and CD27. Preferential Ki-67 expression by fetal T cells indicates a heightened proliferative activity, perhaps influenced by the relatively lymphopenic fetal environment. Reduced proportions of CD45RO^+^ cells among fetal CD4^+^ Tconv and CD8^+^ T cells is consistent with a relative dearth of foreign antigens in utero and corroborates earlier reports of CD45RA expression by most fetal skin CD3^+^ cells^8,9^. Low CD25 and high CD27 expression by fetal cells likely reflects their naïve status^14^. However, CD25 levels might also indicate limiting IL-2 conditions in fetal tissue^25^. Surprisingly, we found that up to 20 percent of CD4^+^ Tconv or CD8^+^ T cells in second trimester fetal skin do express CD45RO. CD45RO^+^ T cells have also been found in the fetal intestines^11^ and constitute up to 5 percent of CD4^+^ T cells in cord blood from healthy term deliveries^6,26^. It is still debated whether the “memory” phenotype of these cells reflects priming to *in utero* antigens^24^ or acquisition of memory markers secondary to cytokine exposure in the lymphopenic fetal environment^27^. Further work is clearly required to define the ontogeny of the CD45RO^+^ CD4^+^ Tconv and CD8^+^ subsets in fetal skin.

To elucidate the distinct functional capacity of fetal versus adult skin Tconv cells, we performed *ex vivo* re-stimulation and found relatively enriched capacity for IFNγ production as compared to IL-13, IL-17A, IL-22. Based on higher CD27 levels among fetal T cells and reports that this TNF superfamily co-stimulatory receptor has a cell-intrinsic capacity to promote Th1 at the expense of Th17 responses^28^, it is tempting to speculate that CD27 expression might influence the functional capacity of fetal skin T cells. IFNγ- and TNFα-producing CD45RO^+^CD4^+^ Tconv cells have also been identified in the fetal intestine, where they are thought to support healthy gut development^5,6^. Notably, numbers of these intestinal “Th1” cells decline before term birth. In contrast, their persistence in preterm infants has been implicated in heightened risk of necrotizing enterocolitis and gastroschisis^5,6^. Premature infants also demonstrate different propensities towards diseases in which skin immune cells play a central role, including increased susceptibility to infections by coagulase-negative staphylococci^29^ and reduced risk of atopic dermatitis (AD)^27-29^. Th17 cells are critical players in the normal response to these commensals^30^. Our finding of reduced IL-17 production by T cells in second trimester fetal skin may be another key factor, alongside skin barrier immaturity, contributing to prematurity-associated infection risk. Conversely, it is intriguing to consider that heightened propensity towards IFNγ in skin may contribute to the lower incidence of AD in preterm infants. Pediatric AD is characterized by Th2 and Th17-mediated inflammation^30,31^, and T cells from the blood of term infants and young children have been shown to display a propensity toward production of type 2 cytokines^37-40^. IFNγ levels in cord blood have also been shown to be conversely associated with atopy at 1 year of age^31^. If early life exposure to microbial and environmental antigens takes place in an IFNγ-rich environment this may steer the response away from an atopic diathesis.

Tregs are relatively well-studied in the fetal immune system but have not been carefully examined in skin. We found Tregs in fetal skin to be quite similar to their adult counterparts in demonstrating an effector memory profile^32^. Modest age-related differences in key markers, however, may indicate distinct functional capacity of skin Tregs by age. As has been shown for Tregs in pediatric tissues such as the spleen and gut^33^, fetal skin Tregs demonstrate higher expression of the master transcriptional regulator Foxp3. PD-1, a co-inhibitory receptor that can mark Tregs with heightened suppressive function as well as stability, was also elevated early^34,35^. Despite being a direct downstream target of Foxp3, CTLA-4 was reduced in fetal versus adult Tregs. Although these data provide some early clues, additional work is merited to tease apart the relative suppressive capacity and regulatory mechanisms used by Tregs in young versus mature human tissues.

Treg enrichment in the fetal immune system is thought to be critical for promoting tolerance to self and maternal antigens^36^ and is supported by multiple mechanisms^30,40,41^. This penchant for tolerance extends beyond the fetal period into early childhood, as indicated by enrichment of Tregs in pediatric versus adult tissues^33,37^ and clinical studies supporting a time-limited early window to promote oral tolerance to food allergens^38^. Here we demonstrate that Tregs progressively increase as a proportion of T cells in fetal skin during the second trimester. More specifically, skin Tregs increase in parallel with terminal hair follicle development, and a relationship between follicle stage and tissue Treg density is preserved at different skin sites within a given fetus. This differs from intestinal Tregs that constitute no more than 10% of CD4^+^ T cells at all fetal time points studied^11^ and suggests that tissue-intrinsic factors, not just progressive thymic output, help mediate Treg accumulation at peripheral sites. Hair follicle-derived chemokines have been implicated in recruitment of various T cell subsets^39^. In neonatal mice, Ccl20 production by hair follicles facilitates postnatal skin Treg accumulation^40^. Analogous chemokine-dependent processes may be active in human fetal skin to recruit recent thymic emigrants. Alternatively, or in addition, an increasing burden of self-antigens contained in terminally-differentiated hair follicles may stimulate local Treg induction or proliferation.

In summary, our findings provide new insight into cutaneous immune development during the second trimester of fetal life. This marks the start of a critical window extending into early childhood during which, specific exposures are thought to influence predisposition to allergic disease. Understanding the skin immune system during this period is of particular relevance given evidence that percutaneous allergen exposure is an early event catalyzing the atopic march. If barriers to tissue access can be surmounted, an important next step will be to define how skin immune cell populations evolve following the transition out of the womb and how this compares between premature and term neonates. Pairing this with investigations to define the immunologic impact of relevant environmental exposures, i.e. commensal microbes or cutaneous allergens, will inform our understanding of healthy development and hopefully better position us to prevent or treat immune-mediated disease in early life.

## Acknowledgements

This work was funded by Doris Duke Clinician-Scientist Development Award 2017070. TCS receives salary support from NIH grant 5K08AR68409 although fetal work was not funded through this mechanism. We appreciate support from lab members of Trevor Burt M.D., Melissa Ng and Ventura Mendoza, who assisted in acquisition of fetal skin samples and UCSF surgeons, Isaac Neuhaus, Scott Hansen, Hobart Harris, and Ester Kim who provided surgically discarded adult skin samples. Flow cytometry and CyTOF data were generated in the UCSF Parnassus Flow Cytometry Core, which is supported by the Diabetes Research Center (DRC) grant, NIH P30 DK063720. We thank Matt Spitzer for input and expertise in development of CyTOF methods for human skin. We appreciate Tim McCalmont’s consultation on histologic examination of fetal hair follicles.

## Author contributions

M.O.D. and T.C.S. drafted the manuscript and figures. M.O.D., K.S.V., E.G.L., K.T., M.M.L. and S.F. performed experiments. M.O.D., M.M.L, M.P, M.D.R. and T.C.S oversaw experimental design. D.K. and M.O.D. performed bioinformatic analysis.

## Materials and Methods

### Human skin specimens

Human fetal tissues (17-23 weeks gestational age) were obtained from Zuckerberg San Francisco General Hospital from terminations of pregnancy after maternal written informed consent with approval from the UCSF Research Protection program. Samples were excluded in the case of (1) known maternal infection, (2) intrauterine fetal demise, and/or (3) known or suspected chromosomal abnormality. Normal adult human skin was obtained from patients at UCSF undergoing elective surgery, in which healthy skin was discarded as a routine procedure. The study was conducted in accordance with the Declaration of Helsinki principles.

### Histology

For histologic analysis of fetal skin, the tissue was fixed in 10% formalin and submitted as research specimens to the UCSF Dermatopathology service, where they were processed, embedded in paraffin, sectioned, and stained with H&E. Hair follicle staging was performed by the investigators in consultation with a dermatopathologist according to previously published criteria ^19^. Images were acquired on a Leica microscope with a DS-Ri1 camera and NIS-Elements software (Nikon).

### Skin digestion

Skin samples were stored at 4°C in a sterile container with PBS and gauze until the time of digestion. Subcutaneous fat was removed, and skin was minced finely with dissection scissors and mixed in a 6-well plate with 3 ml of digestion buffer consisting of 0.8 mg/ml Collagenase Type 4 (4188; Worthington), 0.02 mg/ml DNAse (DN25-1G; Sigma-Aldrich), 10% FBS, 1% HEPES, and 1% penicillin/streptavidin in RPMI medium. Samples were incubated overnight in 5% CO_2_ and harvested with wash buffer (2% FBS, 1% penicillin/streptavidin in RPMI medium), then filtered twice through a 100-μm filter, centrifuged, and counted.

### Flow cytometry

Single cell suspensions were stained for surface antigens and a live/dead marker (Ghost Dye Violet 510, Tonbo Biosciences) in FACS buffer (PBS with 2% fetal bovine serum) for 30 min at 4°C. They were then fixed and permeabilized using the FOXP3-staining buffer kit (eBioscience) before staining for intracellular markers. To measure intracellular cytokine production, cells suspensions were stimulated for 4 hours *ex vivo* using a commercial cell stimulation cocktail (Tonbo Biosciences, catalog no. TNB-4975) prior to staining for flow cytometry as above. Antibodies for the following epitopes were purchased from Biolegend, BD Biosciences, eBiosciences and R&D systems as follows: CD3 (SK7, OKT3), CD4 (S3.5), CD8 (RPA-T8, SK1), CD25 (M-A251), CD27 (LG.7F9), CD31 (WM59), CD45 (2D1), CD45RA (HI100), CD45RO (UCHL1), CD152/CTLA4 (UC10-4B9), CD278/ICOS (ISA-3), Foxp3 (PCH101), IFNγ (4S.B3), IL-13 (85BRD), IL-17A (eBio64CAP17), IL-22 (142928), Ki-67 (B56), TNFα (Mab11). Samples were run on a Fortessa (BD Biosciences) in the UCSF Flow Cytometry Core. FlowJo software (FlowJo LLC) was used to analyze flow cytometry data.

### Mass Cytometry (CyTOF)

Single cell suspensions were incubated for 1 minute with 25 µM Cisplatin (Sigma-Aldrich, P4394) to allow subsequent cell viability measurement, then fixed in 1.5% paraformaldehyde (Electron Microscopy Sciences) and frozen down in the presence of 10% di-methyl sulfoxide (DMSO) and 10% bovine serum albumin (BSA) at stored at −80°C for subsequent staining. Individual vials were thawed and 2×10^6^ cells were barcoded using the Cell-ID 20-Plex Pd Barcoding Kit (Fluidigm, product number 201060) for 15 minutes at room temperature. Barcoded samples were then combined into a single tube and cells were stained with metal conjugated antibodies, purchased either from BD Biosciences or Fluidigm as listed here: CD45 (H130)-89Y, CD19 (HIB19)-142Nd, CD11b (ICRF44)- 144Nd, CD4 (RPA-T4)-145Nd, CD11c (BU15)-147Sm, CD25 (2A3)-149Sm, γδTCR (11F2)-152Sm, CD8α (RPA-T8)-53Eu, CD24 (ML5)-154Sm, CD56 (B159)-155Gd, CD27 (L128)-158Gd, PDL1 (29E.2A3)-159Tb, CD152/CTLA4 (14D3)-14D3, FoxP3 (PCH101)-162Dy, CD1c (L161)- 164Dy, PD1 (EH12.2H7)-166Er, Ki67 (Ki67)-168Er, CD3 (UCHT1)-170Er, CD45RO (UCHL1)-173Yb, HLA-DR (L243)-174Yb, CD207/Langerin (4C7)-175Lu, CD127 (A019D5)-176 Yb. Surface antigens were stained in cell staining media (0.5% BSA, 0.02% Sodium azide in PBS) for 30 minutes at room temperature with gentle agitation. For intracellular staining, cells were permeabilized using the FOXP3-staining buffer kit (eBioscience) and then incubated for another 30 minutes at room temperature with intracellular antibodies in the presence of the FOXP3-staining kit permeabilization buffer. Cells were stored overnight at 4°C in a buffer containing iridium and then run on a Helios CyTOF system (Fluidigm) in the UCSF, Parnassus flow cytometry core facility. FCS files were first processed in FlowJo (FlowJO LLC) to gate on populations of interest, e.g. CD45^+^ (singlet live-CD45^+^), CD4^+^ T cells (CD3^+^CD4^+^CD8^neg^CD56^neg^), CD8^+^ (CD3^+^CD8^+^CD4^neg^CD56^neg^). Further analysis was performed in R using the HDCyto and CATALYST packages found on Bioconductor. Generation of graphs was drawn from the CytofWorkflow pipeline similarly found on Bioconductor ^41^. Multidimensional Scaling (MDS) was used as a first pass at dimensionality reduction and visualization of relatedness of samples as depicted by PCA plot. Uniform Manifold Approximation and Projection (UMAP), an improved method of dimensionality reduction based on greater mathematical rigor than t-SNE, was used to visualize relationships among the single cells on a 2-D plane^42^. Unlike t-SNE, distances between groups on UMAP plots reflect their degree of relatedness capturing the global relations. Clustering into subpopulations, as depicted in Figures 4a-c and S3a, was performed^43^ taking into account all markers in the panel except CD207, CD56, CD1c, CD11c, γδTCR, CD11b, and HLA-DR. Clustering was performed using FlowSOM^44^ and CosnensusClusteringPlus^45^ for metaclustering. Cluster C in the CD4^+^ T cells, identified as Tregs based on their expression profile, was separated from the Teff groups using R scripts to filter data points by cluster.

### Statistical analysis

Significance was determined using either two-tailed unpaired Student’s t test (for measuring differences between separate groups), or one-way analysis of variance (ANOVA) (for multiple comparisons) in GraphPad Prism Software. Data points in all graphs represent individual skin samples. P values correlate with symbols as follows: ns, not significant (P > 0.05); *P < 0.05; **P < 0.01; ***P < 0.001; ****P < 0.0001.

## Supplemental Information

**Figure S1:**
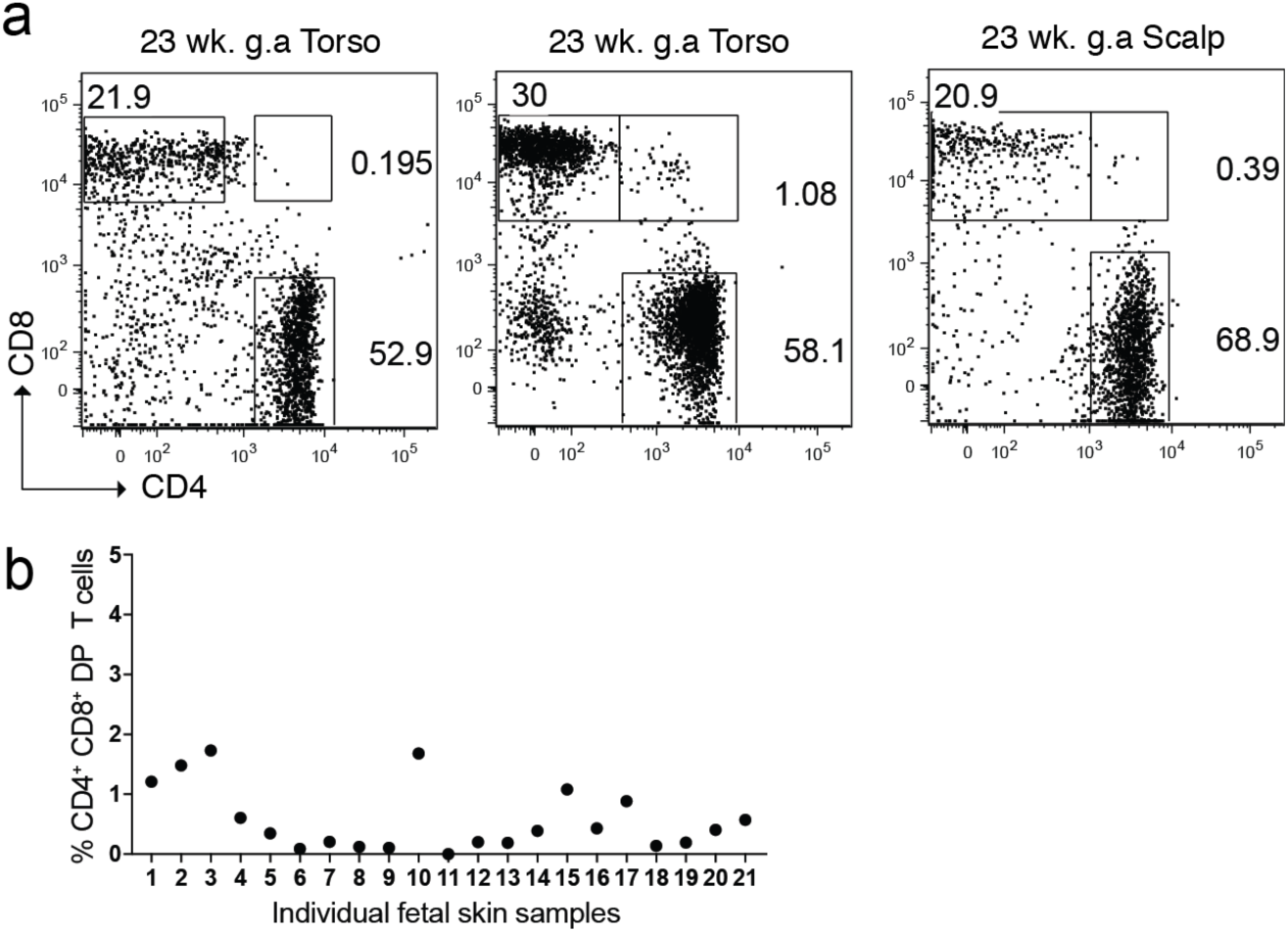
CD4^+^CD8^+^ double-positive T cells constitute a very small population in human fetal skin. Cells were isolated from 23 week g.a. fetal skin (scalp and/or torso) and analyzed by flow cytometry. (a) Representative flow plots of CD4 and CD8 expression by live CD3^+^ lymphocytes in fetal skin. (b) Percentage of DP CD4^+^CD8^+^ lymphocytes across 21 fetal skin samples.

**Figure S2:**
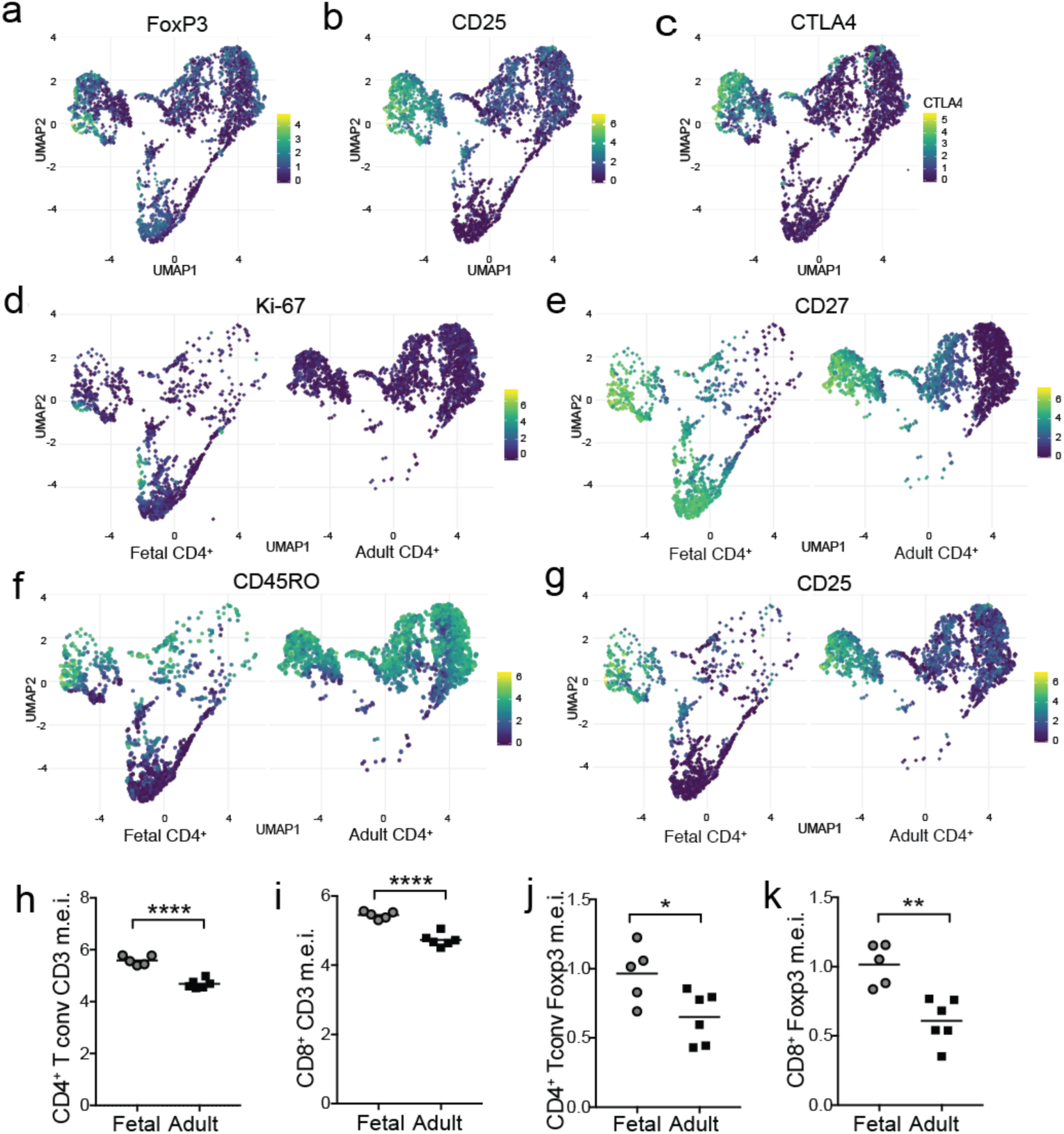
Age-based expression of key markers on skin αβ T cell subsets. 23 week g.a. fetal torso skin along with healthy adult skin torso samples were analyzed in parallel for 22 markers using mass cytometry. (a-c) UMAP plots of single positive CD4^+^ cells from combined fetal and adult skin, colored by intensity of (a) Foxp3, (b) CD25, and (c) CTLA4 expression. (d-e) UMAP plots of single positive CD4^+^ cells from fetal versus adult skin colored by intensity of (d) Ki-67, (e) CD27, (f) CD45RO, and (g) CD25 expression. (h-k) Median expression of CD3 (h-i) and Foxp3 (j-k) on skin CD4^+^ Tconv and CD8^+^ T cells by age. Each point represents data from an individual donor.

**Figure S3:**
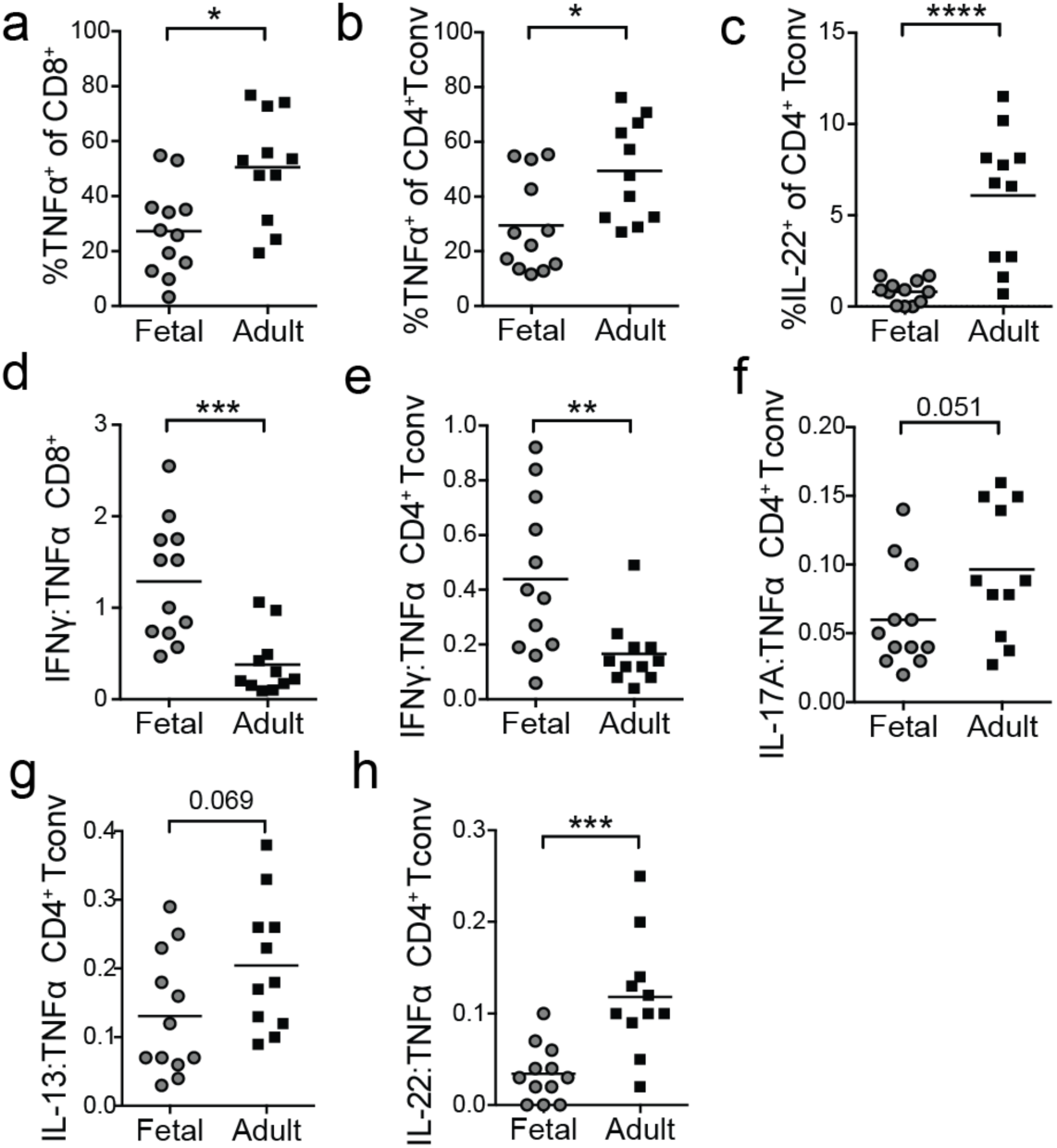
Fetal as compared to adult skin T cells have greater capacity for IFNγ production. Cells were isolated from 23 week g.a. fetal skin (scalp and/or torso) as well as adult (torso) skin and analyzed by flow cytometry following PMA/ionomycin re-stimulation. Percentage TNFα-producing (a) CD8^+^ T cells and (b) CD4^+^ Tconv in skin by age. (c) Percentage IL-22 producing CD4^+^ Tconv by age. (d-h) Cytokine expression was normalized on a per sample basis to the percentage of TNFα^+^ cells. Normalized IFNγ production by (d) CD8^+^ T cells and (e) CD4^+^ Tconv in fetal vs. adult skin. Normalized (f) IL-17A, (g) IL-13 and (h) and IL-22 by CD4^+^ Tconv in fetal vs. adult skin. Each point in represents data from an individual tissue sample; for some fetal samples data from scalp and torso skin from the same fetal donor are included as separate points.

**Figure S4:**
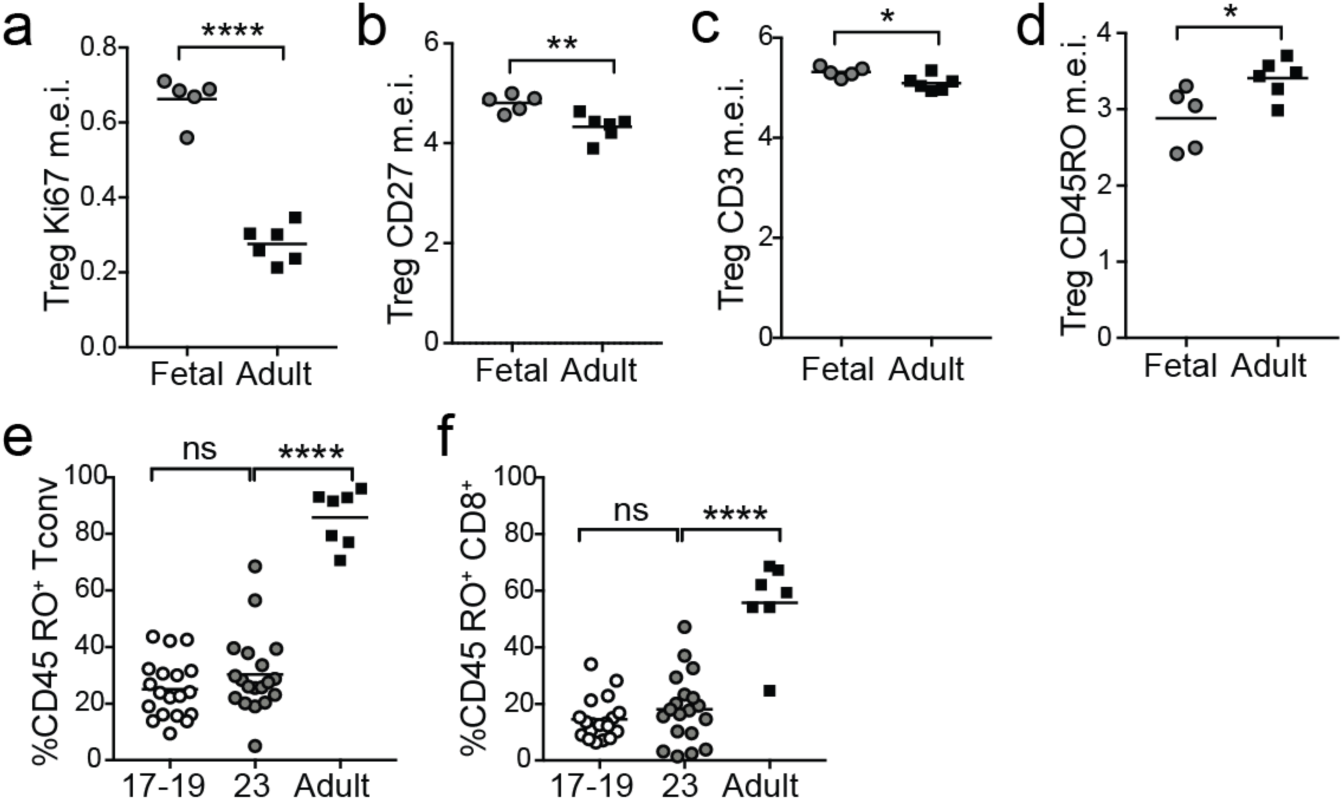
Expression of key markers by fetal vs. adult skin Tregs and of CD45RO by skin CD4^+^ Tconv and CD8^+^ T cells. (a-c) 23 week g.a. fetal torso skin along with healthy adult skin torso samples were analyzed in parallel for 22 markers using mass cytometry. Median expression of (a) Ki-67, (b) CD27, (c) CD3 and (d) CD45RO on skin Tregs by age. Each point represents data from an individual donor. (e-f) Cells were isolated from 17 to 23 week g.a. fetal skin (scalp and/or torso) as well as adult (torso) skin and analyzed by flow cytometry. Percentage of CD45RO^+^ (e) CD4^+^ Tconv cells and (f) CD8^+^ T cells by age. Points represent data from an individual tissue sample; for some fetal samples data from scalp and torso skin from the same fetal donor are included as separate points.

**Figure S5:**
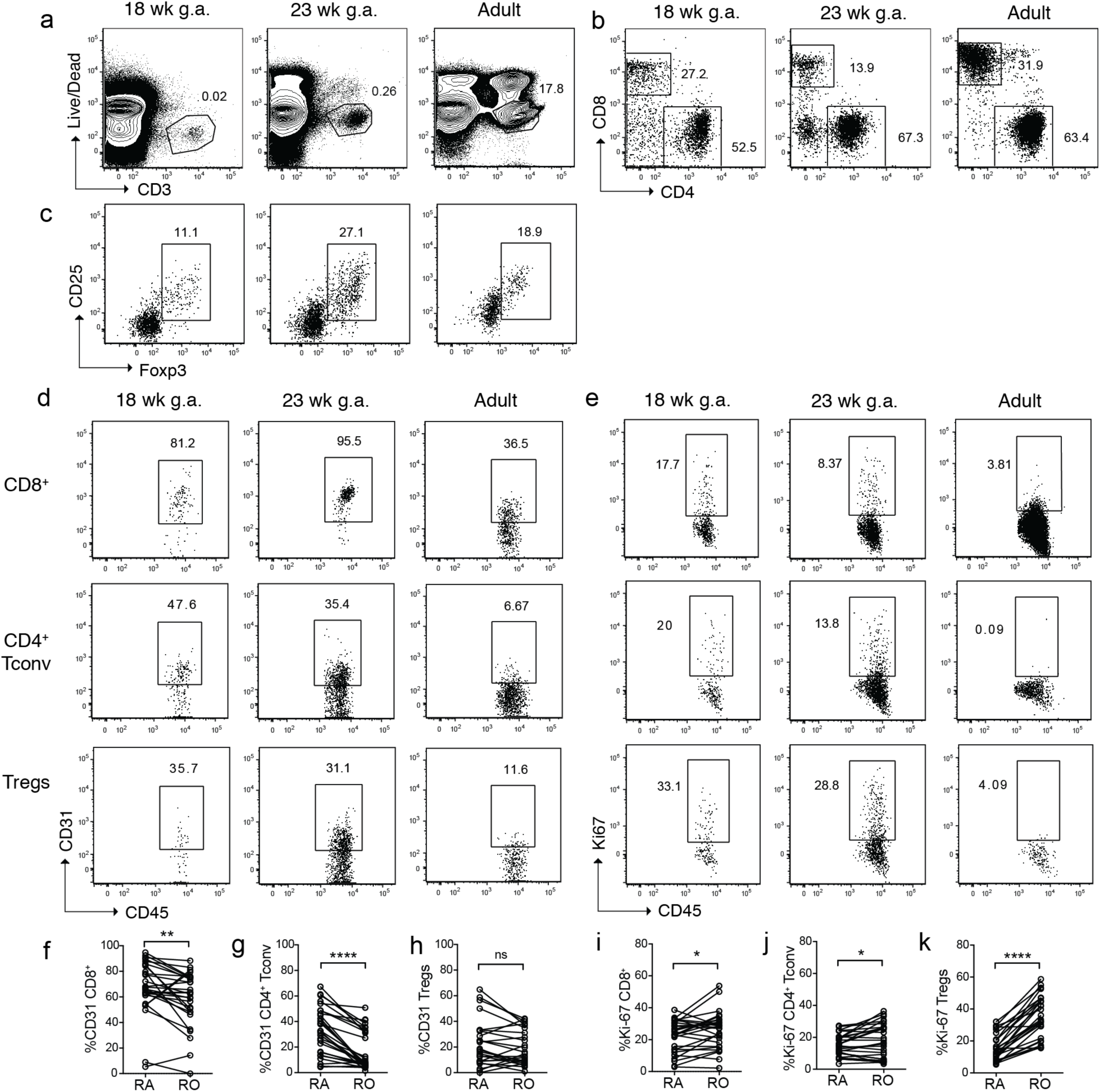
Abundance of αβ T cell subsets and their expression of CD31 and Ki-67 vary by age in human skin. Cells were isolated from 17 to 23 week g.a. fetal skin (scalp and/or torso) as well as adult (torso) skin and analyzed by flow cytometry. Representative flow cytometry plots by age demonstrating (a) live CD3^+^ T cells (pre-gated on singlets), (b) CD4^+^ and CD8^+^ expression by T cells (pre-gated on live CD3^+^) and (c) Tregs (pre-gated CD3^+^CD8^neg^CD4^+^). Representative flow plots by age showing (d) CD31 and (e) Ki-67 expression by skin αβ T cell subsets. Percentage of CD31^+^ cells among CD45RA^+^ vs. CD45RO^+^ (f) CD8^+^ T cells, (g) CD4^+ -^Tconv cells, and (h) Tregs in fetal skin. Percentage of Ki-67^+^ cells among CD45RA^+^ vs. CD45RO^+^ (i) CD8^+^ T cells, (j) CD4^+ -^Tconv cells, and (k) Tregs in fetal skin. (f-k) Paired data points represent CD45RA^+^ vs CD45RO^+^ subsets from the same tissue sample.

